# Postpartum breast cancer progression is driven by semaphorin 7a mediated invasion and survival

**DOI:** 10.1101/631044

**Authors:** Sarah E Tarullo, Ryan C Hill, Kirk Hansen, Fariba Behbod, Virginia F Borges, Andrew C Nelson, Traci R Lyons

**Affiliations:** Department of Medicine, Division of Medical Oncology, CU Anschutz Medical Campus, Aurora, CO 80045; Young Women’s BC Translational Program, CU Anschutz Medical Campus, Aurora, CO 80045; Department of Biochemistry and Molecular Genetics, CU Anschutz Medical Campus, Aurora, CO 80045; Division of Cancer and Developmental Biology, University of Kansas Medical Center, Kansas City, KS 66160; University of Colorado Cancer Center, Aurora, CO 80045; Department of Laboratory Medicine and Pathology, University of Minnesota, Minneapolis, MN 55455

## Abstract

Young women diagnosed with breast cancer (BC) have poor prognosis due to increased rates of metastasis. Additionally, women within 10 years of most recent childbirth at diagnosis are ∼3 times more likely to develop metastasis than age and stage matched nulliparous women. We define these cases as postpartum BC (PPBC) and propose that the unique biology of the postpartum mammary gland drives tumor progression. Our published results revealed roles for SEMA7A in breast tumor cell growth, motility, invasion, and tumor associated-lymphangiogenesis, all of which are also increased in pre-clinical models of PPBC. However, whether SEMA7A drives progression in PPBC remains largely unexplored. Our results presented herein show that silencing of SEMA7A decreases tumor growth in a model of PPBC while overexpression is sufficient to increase growth in nulliparous hosts. Further, we show that SEMA7A promotes multiple known drivers of PPBC progression including tumor associated COX-2 expression and fibroblast-mediated collagen deposition in the tumor microenvironment. Additionally, we show for the first time that SEMA7A expressing cells deposit fibronectin to promote tumor cell survival. Finally, we show that co-expression of SEMA7A/COX-2/FN predicts for poor prognosis in breast cancer patient cohorts. These studies suggest SEMA7A as a key mediator of BC progression and that targeting SEMA7A may open avenues for novel therapeutic strategies.

## INTRODUCTION

Postpartum breast cancer (PPBC), or breast cancers (BC) diagnosed within 5-10 years of last childbirth, are ∼three times more likely to become metastatic [1–3]. Specifically, PPBC patients exhibit distant metastasis free five-year survival (DMFS) rates as low as 70% [1], which are further decreased to 50% after ten years [2]. Additionally, PPBC may account over half of BCs diagnosed in women aged <45 [1]. In a pre-clinical model of PPBC a non-metastatic BC cell line becomes invasive and metastatic upon orthotopic implantation at the onset of postpartum mammary gland involution [4]. Postpartum/post-lactational mammary involution returns the gland to the pre-pregnant state; we and others have shown that programs associated with postpartum involution are similar to tumor-promotional microenvironments [1, 4–9]. Since the MCF10DCIS model initially resembles ductal carcinoma in situ (DCIS), which progresses to ER/PR/HER2 negative invasive ductal carcinoma (IDC) [10], we utilized this model to show accelerated tumor growth and progression to IDC in postpartum hosts [4]. This progression was driven by collagen deposition and expression of cyclooxygenase-2 (COX-2), both of which were required for tumor cell invasion. Additionally, a weakly tumorigenic breast epithelial cell line, HMLE-Ras^lo^, was similarly promoted via host driven mechanisms [11]. More recently, expression of a neuronal guidance molecule, Semaphorin 7a (SEMA7A), was observed in mouse mammary epithelium during postpartum involution and in the tumors that outgrew after implantation during involution [6].

Semaphorins are characterized for their roles during development, however and have reported roles in multiple cancer types [12–16] and SEMA7A expression is emerging as poor prognostic indicator [6, 16–20]. SEMA7A can promote cell-autonomous signaling when it remains bound to the cell via its glycosylphosphatidylinositol (GPI) membrane link or non-cell-autonomous signaling when shed via cleavage into the extracellular environment. Here, we show that SEMA7A protein is expressed in DCIS from BC patients, is necessary for postpartum tumor progression in our pre-clinical model, and sufficient to drive tumor progression in nulliparous hosts. We also demonstrate that shed SEMA7A drives collagen deposition in the tumor microenvironment (TME) via upregulation of collagen I mRNA in fibroblasts, which promotes expression of COX-2 and invasion. Furthermore, we propose a cell-autonomous pro-invasive and survival role for SEMA7A that is mediated through fibronectin (FN), epithelial-to-mesenchymal transition (EMT) and downstream pro-survival signaling via phosphorylation of AKT. Additionally, we show that SEMA7A expressing cells exhibit enhanced metastatic capabilities. Finally, our results we demonstrate that a gene signature of SEMA7A, COX-2, and FN1 predicts for poor prognosis for BC patients suggest that SEMA7A merits further studies to develop a novel therapeutic for BC patients.

## RESULTS

### SEMA7A promotes postpartum tumor progression in a mouse model and is expressed in DCIS from patients

The MCF10DCIS model is ideal for monitoring early events in the metastatic cascade, such as the transition from in situ to invasive. To examine whether SEMA7A drives DCIS progression in postpartum/post-lactational hosts, we orthotopically injected MCF10DCIS cells stably expressing a SEMA7A targeted shRNA (SEMA7A-KD) or nontargeting control (Crtl) [18] at involution day 1 (SFigure1A). We observed that silencing of SEMA7A is sufficient to decrease tumor growth in postpartum hosts despite some of the tumors regaining expression of SEMA7A protein (Figure1A; SFigure1B). Harvested tumors (H&E stained sections) were scored for invasion at 5 weeks post-injection, when the majority of postpartum tumors are normally invasive (SFigure1C), and 33% of the SEMA7A-KD were invasive compared to 86% in the controls. Further, 56% of tumors in the SEMA7A-KD group maintained evidence of DCIS versus 12.5% controls (Figure1B). Finally, 11% of tumors in the KD group were DCIS with microinvasion compared to 0% of controls. Collagen-mediated upregulation of COX-2 is a dominant feature that drives invasion in postpartum hosts in the MCF10DCIS model [4]. Consistent with a role for SEMA7A in collagen/COX-2 dependent invasion in postpartum hosts, we observe that fibrillar collagen deposition, by Massan’s trichrome stain, and COX-2 expression, by immunohistochemistry (IHC) are significantly decreased in SEMA7A-KD tumors (Figure1C&D).

**Figure 1.**
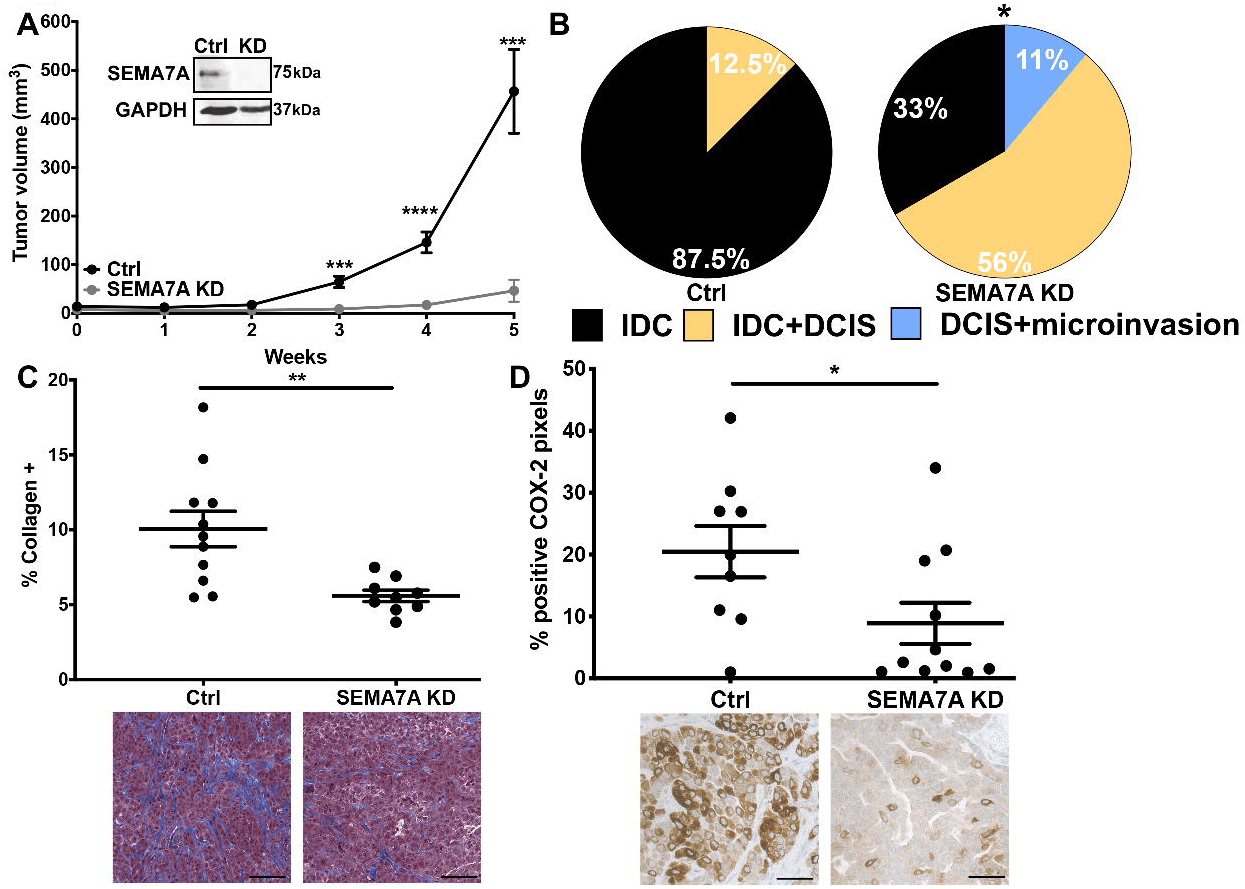
SEMA7A promotes growth and invasion in PPBC. **A.** Tumor volumes for control (Ctrl) or shSEMA7A (SEMA7A-KD) MCF10DCIS cells in postpartum hosts (n=12/group), inset: representative immunoblot for SEMA7A and GAPDH. **B.** Tumors from Ctrl and SEMA7A-KD from postpartum hosts scored for invasion (n=12/group). **C**. Trichrome stained quantification (top) and representative images (bottom) of collagen of tumors from Ctrl and SEMA7A-KD tumors from postpartum hosts, scale bars 50μm (n=12/group) . **D.** IHC quantification (top) and representative images (bottom) of IHC for COX-2 of tumors from Ctrl and SEMA7A-KD tumors from postpartum hosts, scale bars 50μm (n=12/group) (*p<0.05, **p<0.01, ***p<0.0005, ****p<0.0001, t-test).

To determine whether SEMA7A expression was higher in DCIS from postpartum patients, we performed IHC on DCIS lesions and adjacent normal breast tissue from women in our BC cohort (Figure 2A) and quantitated medium + strong staining to show that DCIS lesions express higher levels of SEMA7A than normal breast tissue (Figure2B,SFigure2A). Then, when separated by parity status, we observe increased SEMA7A in DCIS lesions in both nulliparous and postpartum patients, with a trend toward highest expression in postpartum patients, suggesting that SEMA7A expression may have relevance to all patients with DCIS, but may be particularly relevant to postpartum patients (Figure2B,SFigure2B). Consistent with this, paired analysis between normal and DCIS within each patient revealed that the majority of patients in both the nulliparous and postpartum groups exhibited increased SEMA7A expression in DCIS compared to normal (Figure 2C&D). We confirmed this observation by showing similar data on SEMA7A expression in a tissue microarray consisting of additional normal breast and matched patient samples of DCIS using kidney and placenta as negative and positive controls, respectively (Figure2E-G,SFigure2C). Finally, we also show that SEMA7A mRNA levels are increased in DCIS compared to normal in the METABRIC [21] dataset and that SEMA7A is in the top 11% of upregulated genes in DCIS compared normal (p=0.002)(SFigure2D). Taken together, our results suggest that SEMA7A may represent a general mediator of DCIS growth and invasion in both parous and nulliparous women.

**Figure 2.**
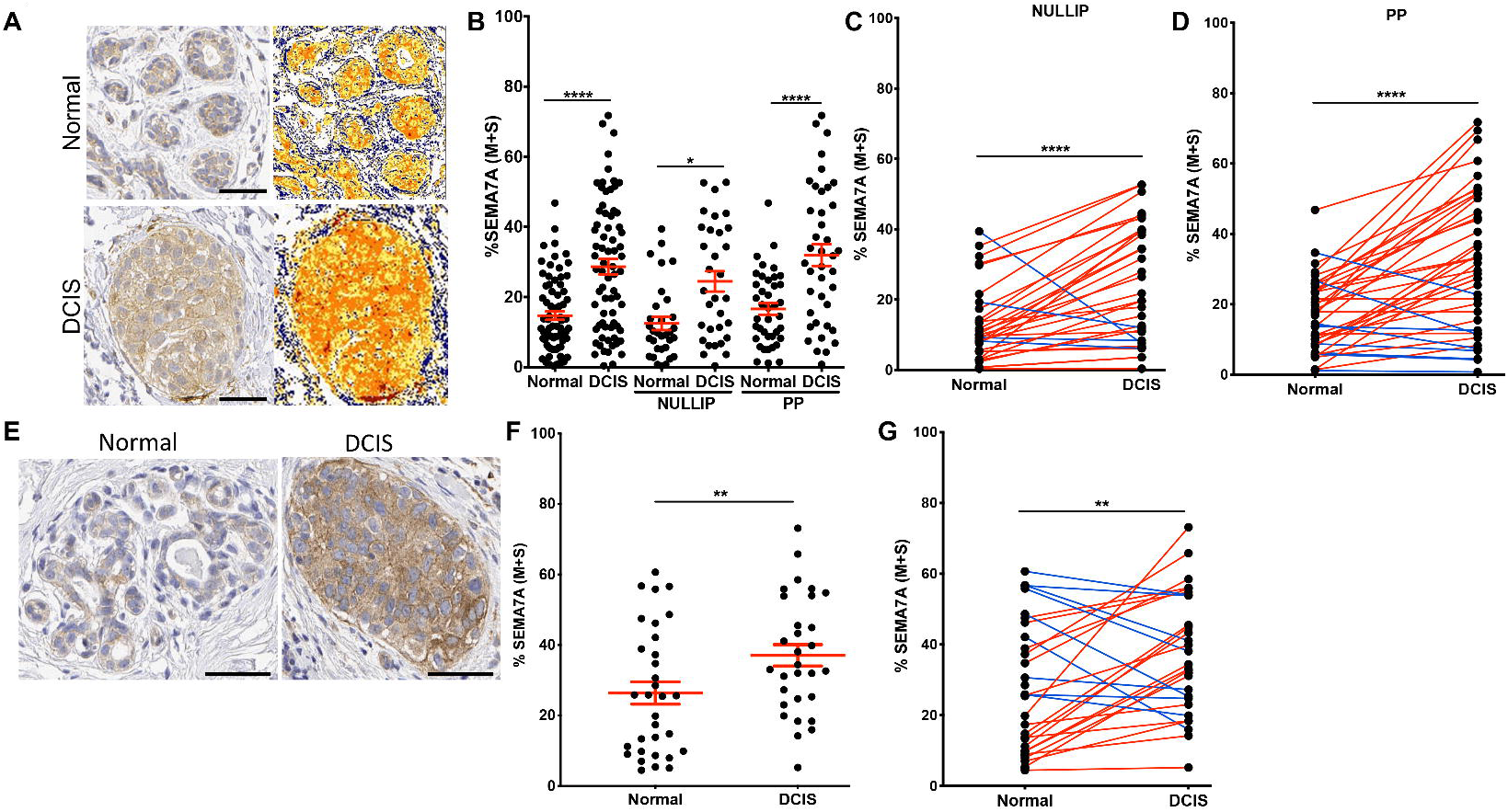
SEMA7A expression is increased in DCIS patient samples. **A.** Representative images of normal (top) or DCIS lesions (bottom) from postpartum patients from the CU Anschutz Medical Campus Young Women’s BC (YWBC) stained by IHC for SEMA7A, with quantification mask to the right, blue=negative; yellow=weak; orange=medium; red=strong; scale bars 50 μm. **B.** Quantification of SEMA7A IHC of normal or DCIS patient samples from the CU Anschutz YWBC cohort (n=103), then separated by parity status, nulliparous (n=52) or postpartum (n=51). **C**. Quantification of SEMA7A IHC from nulliparous patients by patient, paired t-test. **D.** Quantification of SEMA7A IHC from postpartum patients by patient, paired t-test. **E.** Representative images of SEMA7A IHC stain from tissue array of samples, scale bars 50μm. **F.** Quantification of SEMA7A IHC stain from **C**; normal (n=32) or DCIS (n=30). **G.** Quantification of SEMA7A IHC by patient, paired t-test. (*p<0.05, **p<0.01, ****p<0.001)

To test our hypothesis that SEMA7A expression is sufficient to drive DCIS progression, we injected MCF10DCIS cells that overexpress SEMA7A protein (SEMA7A-OE), along with controls, into a separate cohort of nulliparous hosts; we observed accelerated growth of SEMA7A-OE tumors (Figure3A). These tumors were similarly scored for invasion, but at three weeks post-injection, when tumors are normally DCIS in nulliparous hosts (SFigure1C). While, control tumors were all DCIS, as expected, greater than 50% of the SEMA7A-OE tumors had microinvasion and/or were mixed IDC+DCIS (Figure3B;SFigure 3). Additionally, levels of collagen and COX-2, which are normally very low in DCIS tumors in nulliparous hosts, are higher in the SEMA7A-OE tumors, which is consistent with a role for SEMA7A in promoting production (Figure3C&D).

**Figure 3.**
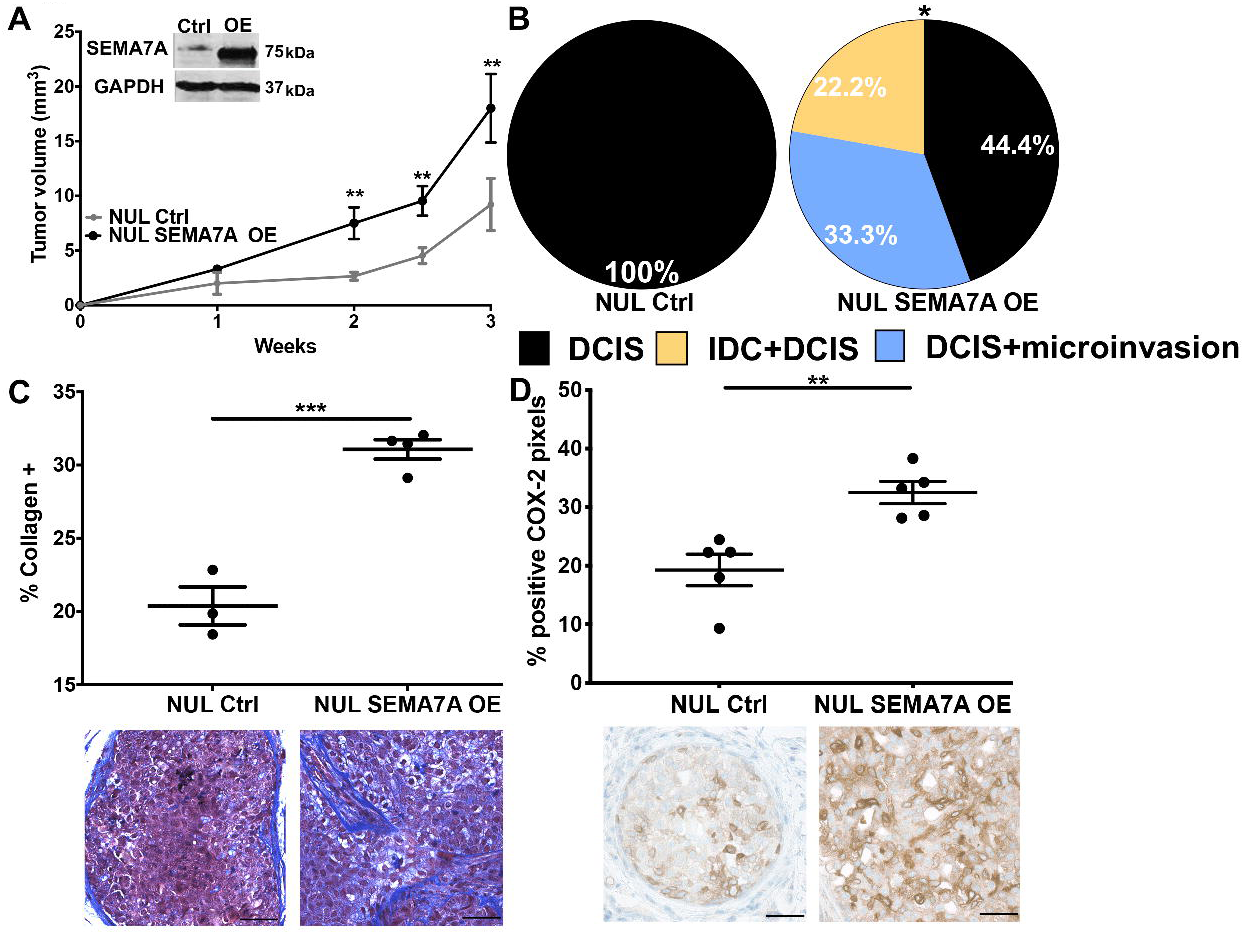
SEMA7A expression is sufficient to drive tumor growth and invasion. **A.** Tumor volumes for control (NUL Ctrl) or SEMA7A overexpressing (NUL SEMA7A OE) in nulliparous hosts (n=10/group), inset: representative immunoblot for SEMA7A and GAPDH. **B**. Tumors from NUL Ctrl and NUL SEMA7A OE scored for invasion (n=10/group). **C**. Trichrome stained quantification (top) and representative images (bottom) of collagen of tumors from NUL Ctrl and NUL SEMA7A OE tumors, scale bars 50μm **D.** IHC quantification (top) and representative images (bottom) of IHC for COX-2 of tumors from NUL Ctrl and NUL SEMA7A OE tumors, scale bars 50μm (*p<0.05, **p<0.01, ***p<0.0005, ****p<0.0001, t-test).

### SEMA7A promotes tumor cell invasion via matrix deposition and acquisition of mesenchymal phenotypes

To model SEMA7A dependent invasion in vitro, we utilized a 3D organoid model with SEMA7A-OE cells suspended in Matrigel or Matrigel+collagen [4]. We found that SEMA7A overexpression was not sufficient to drive increased invasion on Matrigel alone but sufficient when collagen I was present in the matrix (Figure4A&B). We then validated the requirement for both collagen and SEMA7A in invasion via knockdown in the MDA-MB-231 cell line, which requires collagen for 3D organoid formation [22] (SFigure4A, Figure4C). Since fibroblasts in the TME produce collagen, we then stained for alpha-smooth muscle actin (*α*SMA), a marker of activated fibroblasts, but did not observe differences in fibroblast infiltration (SFigure4). These results suggest that SEMA7A does not recruit tumor infiltrating fibroblasts.

**Figure 4.**
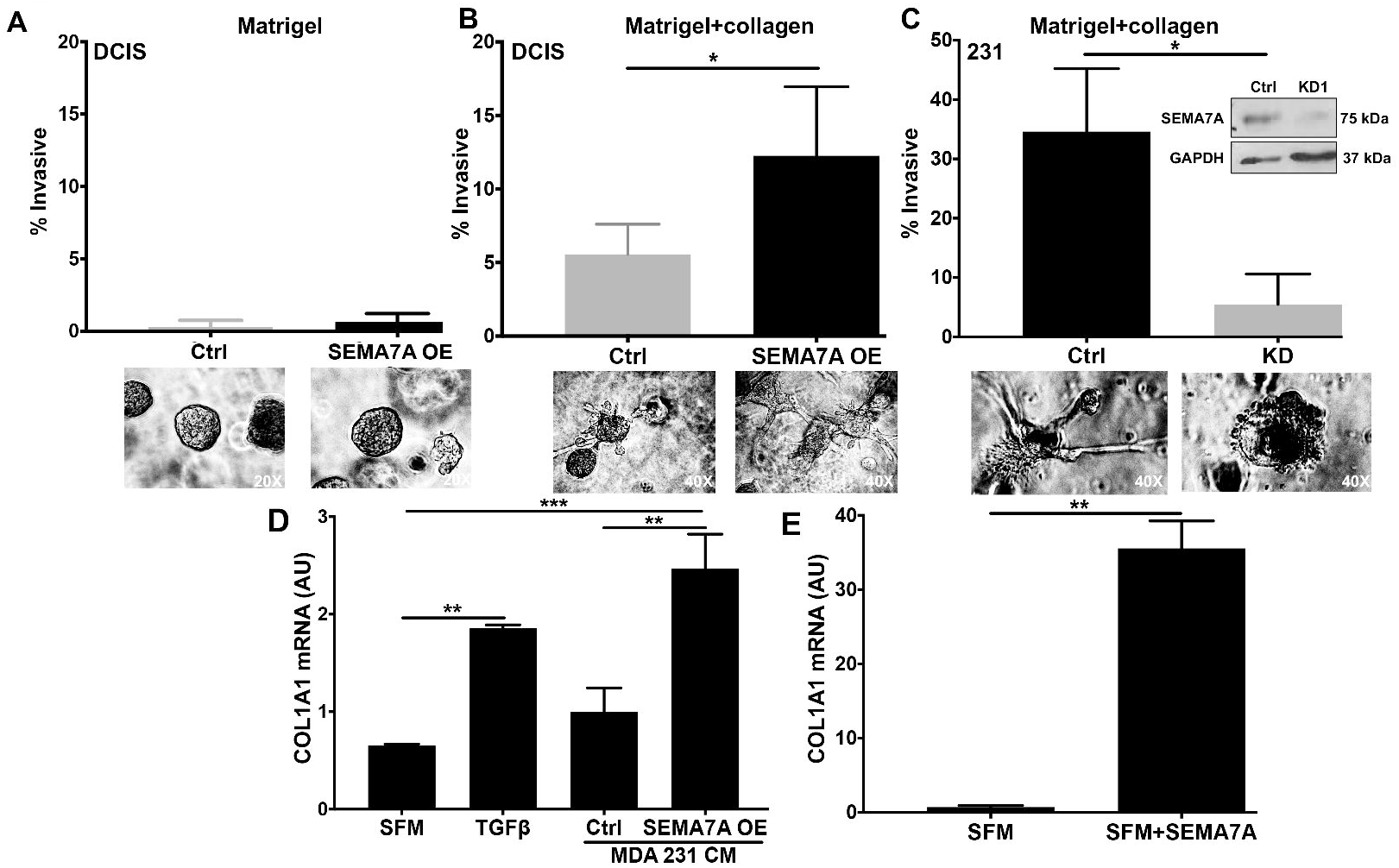
SEMA7A promotes invasion via fibroblast mediated collagen deposition. **A.** Ctrl or SEMA7A OE MCF10DCIS cells embedded in matrigel scored for invasion. **B.** Ctrl or SEMA7A OE MCF10DCIS cells embedded in matrigel plus 20% collagen. **C**. Control (Ctrl) or shSEMA7A (KD) MDA-MB-231 cells in matrigel plus 25% collagen scored for invasion. **D**. Quantitative RT-PCR (q-PCR) for COL1A1 in fibroblasts in serum free media (SFM) treated with 10 ng/mL TGFβ or conditioned media (CM) from control (Ctrl) or SEMA7A overexpressing (SEMA7A OE) MDA-MB-231 cell lines. **E**. qPCR for COLA1A in fibroblasts in SFM treated with 75 ng/μL purified SEMA7A. (*p<0.05, **p<0.01,***p<0.005, t-test).

Alternatively, to determine whether SEMA7A expressing tumor cells promote collagen production by fibroblasts via shedding of SEMA7A, we induced serum starved fibroblasts with conditioned media from our SEMA7A-OE tumor cells, alongside conditioned media from control cells and TGFβ as positive control and examined expression of the *COL1A1* mRNA by qPCR. We focused on *COL1A1* because collagen I is the most abundant collagen in the mammary gland [23]. Conditioned medias from SEMA7A OE cells induced *COL1A1* expression in fibroblasts to higher levels than observed in control or TGFβ treated fibroblasts (Figure4D). Then, we treated fibroblasts with purified SEMA7A to show specific induction of COL1A1 gene expression (Figure4E). To understand additional SEMA7A mediated changes to the TME, we performed an unbiased mass spectrometry analysis of conditioned medias from ex vivo tumors derived from SEMA7A-KD or control cells. When we restricted our analysis to ECM peptides of human origin we observed downregulation of several ECM associated molecules with SEMA7A-KD (SFigure4A), but focused on the significant decrease in fibronectin (FN) (Figure5A) after immunostaining of our tumors confirmed decreased FN expression with SEMA7A-KD (Figure5B&C). As FN is frequently associated with EMT, we examined whether SEMA7A expression is also associated with additional mesenchymal markers. We observed SEMA7A dependent expression of matrix remodeling enzyme, matrix metalloproteinase-2, vimentin, and S100A4, as well as other members of the S100 family of proteins (Figure5D-F;SFigure4B-E), which are all secreted by mesenchymal-like cells [24]. To confirm SEMA7A-dependent mesenchymal-like phenotypes we measured cell aspect ratios with the prediction that epithelial-like cells, due to their cuboidal morphology, would exhibit a ratio of ∼1 and ratios >1 would be indicative of mesenchymal-like cells. Our results reveal that the mesenchymal-like MDA-MB-231 cells, exhibit decreased average aspect ratios with SEMA7A-KD (Figure5G). Concordantly, SEMA7A-OE in the MCF10DCIS cells, which are less mesenchymal-like, resulted in increased aspect ratios (Figure5H). Furthermore, immunoblot analysis of MCF10DCIS SEMA7A-OE cells reveals decreased E-cadherin and increased vimentin expression, markers of epithelial and mesenchymal cells, respectively (SFigure5F). Together, our results suggest that SEMA7A promotes cellular invasion via both cell-autonomous and non-cell-autonomous mechanisms of matrix deposition and remodeling.

**Figure 5.**
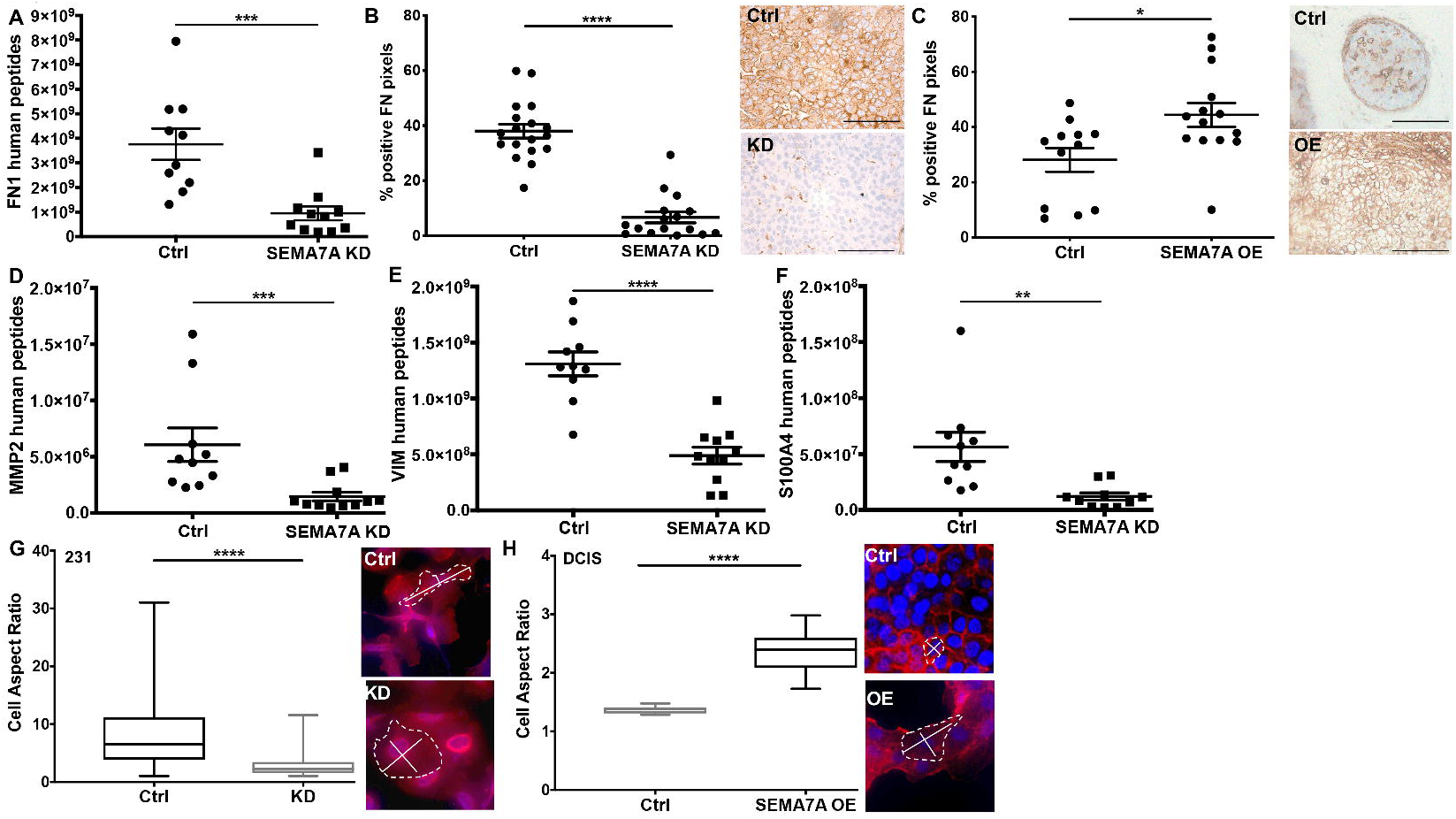
SEMA7A promotes mesenchymal protein expression and phenotypes. **A.** Global proteomics analysis of secreted proteins reveals decreased fibronectin (FN) in MCF10DCIS shSEMA7A (SEMA7A-KD) tumors compared to control (Ctrl) ex vivo (n=10-11/group). **B&C**. IHC for FN in MCF10DCIS Ctrl and SEMA7A-KD tumors (B), and MCF10DCIS Ctrl and SEMA7A overexpressing (SEMA7A OE) tumors, scale bar 50μm. **D-F**. Additional decreases in mesenchymal or mesenchymal associated proteins observed by proteomics in shSEMA7A (SEMA7A-KD) tumors ex vivo. **G**. Immunofluorescence for F-actin in MDA-MB-231 Ctrl or KD cells and quantification for cell aspect ratio (length/width). **H**. Immunofluorescence for F-actin in MCF10DCIS Ctrl or SEMA7A OE cells and quantification for cell aspect ratio (length/width). (*p<0.05, **p<0.01, ***p<0.005, ****p<0.001, t-test)

### SEMA7A promotes cell survival via fibronectin, AKT and COX-2

We also observed decreased tumor growth of SEMA7A-KD tumors in vivo but did not see significant changes in proliferation marker Ki67 (SFigure5A). However, we did observe a trend toward increased cleaved-caspase 3, a marker of apoptosis, in SEMA7A-KD tumors with a corresponding decrease in SEMA7A-OE tumors (SFigure5B&C). We also observed increased cell death in SEMA7A-KD cells in culture via a luminescent assay for caspase activity, which we validated in the MDA-MB-231 cell line (Figure6A&B). Conversely, decreased cell death was observed in the SEMA7A OE cell in both cell lines (SFigure5D, SFigure5E&F). Additionally, analysis of cell death in real time confirmed that SEMA7A-KD MDA-MB-231 cells exhibit increases in cell death via activation of apoptotic signaling (Figure6C). One known pro-survival mechanism co-opted by tumors to block activation of caspase cleavage is activation of pro-survival kinase AKT [25] and we observed decreased levels of phosphorylated AKT via immunoblot for pS473 in SEMA7A-KD and a corresponding increase in levels in SEMA7A OE cells (Figure6D&SFigure5G). Interestingly, FN signals via integrins, which can activate downstream pro-survival pathways, including AKT [26]. To determine whether add-back of FN could rescue SEMA7A-KD cells from cell death, we KD plated cells FN-, laminin- or collagen-coated plates with control cells on tissue culture plastic for reference. We observed increased cell death in all SEMA7A-KD conditions except when FN was present where we observed levels of cell death were more similar to the Ctrl cells (Figure6E). Thus, we suggest that SEMA7A promotes AKT mediated survival by increasing FN deposition and therefore cellular attachment.

**Figure 6.**
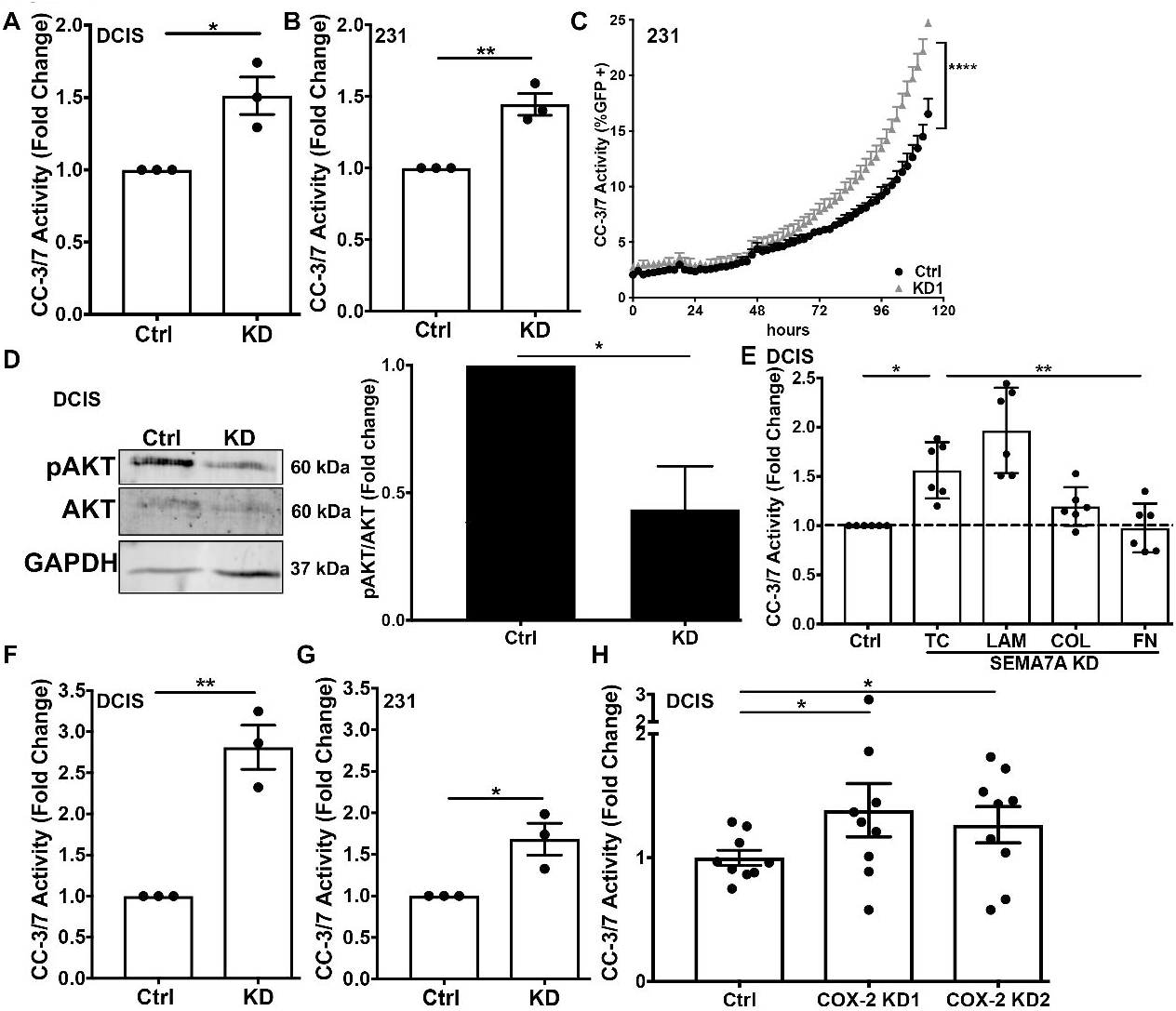
SEMA7A promotes cell survival via fibronectin. **A&B.** Fold change of cleaved caspase 3/7 activity in control (Ctrl) and shSEMA7A (KD) in MCF10DCIS (**A**) or MDA-MB-231 (**B**) cell lines. **C.** Cleaved caspase 3/7 activity measured over time in MDA-MB-231 Ctrl or KD cells. **D.** Representative immunoblot for phospho-AKT (S473), total AKT, or GAPDH in Ctrl or KD, quantified to the right. **E**. Cleaved caspase 3/7 activity control (Ctrl) or shSEMA7A (KD) cells plated on tissue culture plastic (TC), laminin (LAM), collagen I (COL) or fibronectin (FN). **F&G.** Cleaved caspase 3/7 activity MCF10DCIS (**F**) or MDA-MB-231 (**G**) Ctrl or KD cells in forced suspension. **H.** Cleaved caspase 3/7 activity in MCF10DCIS Ctrl or shCOX-2 (KD1/KD2) cell lines in forced suspension.

Mesenchymal-like tumor cells, known to have high levels of FN, are not as dependent on matrix attachment for survival and can also exhibit resistance to normal programs that mediate cell death including survival in detached or anchorage independent conditions [27]. An initial step of the metastatic cascade involves cell detachment from ECM to facilitate local invasion and access to the vasculature; then, cells in circulation must survive in matrix detached conditions prior to extravasating to seed metastatic sites. To assess a role for SEMA7A in promoting survival in circulation, we forced cellular detachment in vitro and measured cleaved-caspase 3/7 to show that SEMA7A-KD increases cell death while SEMA7A-OE promotes cell survival in detached conditions (Figure6F&G, SFigure5H&I). Interestingly, fibronectin was not sufficient to rescue cell death in suspension suggesting that fibronectin deposition is necessary (data not shown). However, since activation of AKT is a known mediator of COX-2 expression [28] we also examined the ability of COX-2 KD cells to survive in detached conditions. Similar to our results in the SEMA7A-KD cells we observe that COX-2 knockdown cells exhibit increased cell death in detached conditions, which was not observed in attached conditions (SFigure5J) suggesting that SEMA7A mediated upregulation of AKT signaling, and subsequent expression of COX-2, support cell survival in detached conditions such as those encountered during local invasion and in circulation (Figure 6H).

### SEMA7A, COX-2, FN and metastatic potential

To assess whether SEMA7A promotes survival in circulation in vivo, we performed tail vein injections of MDA-MB-231 SEMA7A-KD or Ctrl cells (SFigure6A) and assessed for pulmonary metastasis. Our results reveal a decrease in both the number and average size of metastatic lesions per lung with SEMA7A KD (Figure7A) suggesting that SEMA7A plays a role in survival in circulation, as well as a possible role in seeding and outgrowth of BC metastasis. We have previously published that SEMA7A expression is upregulated in IDC compared to normal and the worst overall survival (OS) was observed in SEMA7A+ER-BC in the METABRIC dataset [18]; however, in SEMA7A+ER-BCs we do not observe decreased distant metastasis free survival (DMFS) via KmPlot analysis (SFigure6B). Similarly, although transcripts for COX-2 and FN1 are also upregulated in BC (STable1), neither COX-2 or FN1 are associated with decreased DMFS in ER-BC (SFigure6E&F). However, co-expression of COX-2, SEMA7A, and FN1 increases risk for metastasis in ER-BC patients, with 5 year DMFS rates approaching 70% by both KmPlot and GOBO (Gene expression-based Outcome for Breast cancer Online or GOBO) analysis (Figure7B&C) [1]. We also observed that co-expression of our 3 gene signature significantly associates with decreased DMFS in basal and PAM50 basal subtype tumors using GOBO (SFigure6G-I). Finally, co-expression of SEMA7A with both FN and COX-2 is observed in BCs in the TCGA dataset (Figure6D&E).

**Figure 7.**
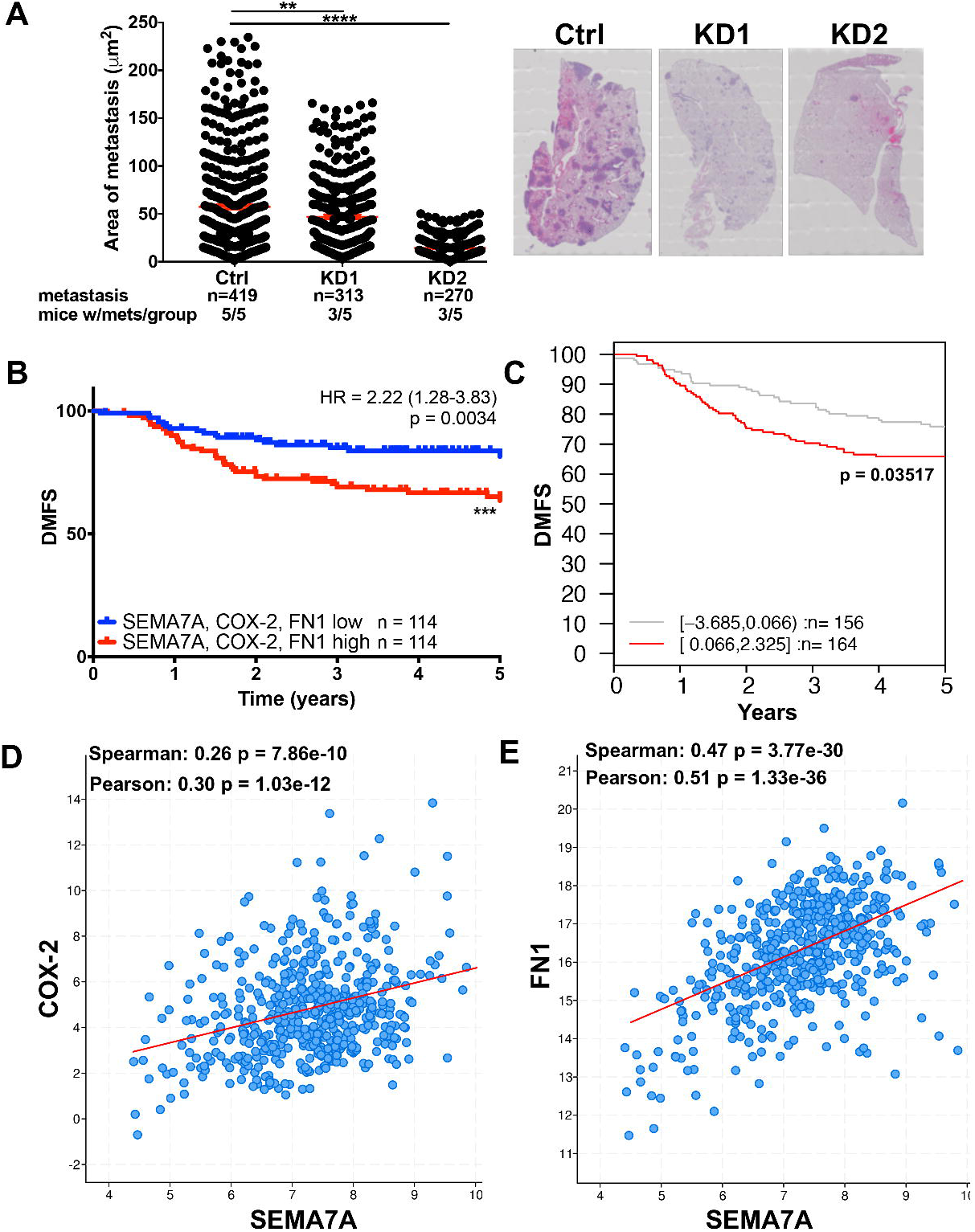
SEMA7A drives metastatic seeding and poor prognosis in patients. **A** Average area of metastasis in lungs of mice after tail vein injection with MDA-MB-231 control (Ctrl) or shSEMA7A (KD1, KD2) cells (n=5/group), representative images of lungs to the right. **B.** Kaplan-Meier analysis of ER-BCs using Km plotter for SEMA7A, COX-2 and FN mRNA expression for distant metastasis free survival (DMFS) (n=228). **C**. Kaplan-Meier analysis of ER-BCs in the Ma Breast Cancer dataset using GOBO for SEMA7A, COX-2 and FN mRNA expression for distant metastasis free survival (DMFS) (n=320) **D&E.** Co-expression analysis of the TCGA BC provisional cohort using CBioPortal for all BCs (n=1108). (*p<0.05, **p<0.01,***p<0.005 ****p<0.001).

## DISCUSSION

All women experience a transient increase in risk for developing breast cancer after each completed pregnancy [4, 29, 30]. Additionally, patients diagnosed with breast cancer within ten years postpartum comprise nearly 50% of all BCs in women <40 in both US and Norwegian cohorts and these PPBC patients are at high risk for metastatic spread [1, 31]. A recent publication by Welch and Hurst defines the hallmarks of metastasis as motility and invasion, modulation of the microenvironment, plasticity, and colonization [32]. Here, we identify roles for SEMA7A in promotion of metastasis using preclinical models regardless of parity status. Our results suggest that SEMA7A drives one described mechanism of tumor progression pertinent to postpartum women, which is collagen mediated upregulation of COX-2 that drives motility and invasion of DCIS cells. However, our results also suggest that this mechanism of invasion can be driven by SEMA7A in both nulliparous and postpartum hosts suggesting it is not unique to parous patients. Further, we extend our observations and identify a novel cell-autonomous mechanism by which SEMA7A mediates tumor cell survival via stimulation of FN production and activation of pro-survival kinase AKT regardless of the host parity status. We also show that SEMA7A supports tumor cell invasion by stably altering cells to a more mesenchymal phenotype, which promotes cell survival in anchorage-independent conditions to allow for colonization of distant organs. Thus, we suggest that SEMA7A plays a key role in multiple steps toward progression to metastatic disease. We support this claim by showing a that co-expression of COX-2, SEMA7A, and FN correlates with distant metastasis formation in BC patients. We propose a model whereby SEMA7A signaling supports metastatic progression (Figure8).

**Figure 8.**
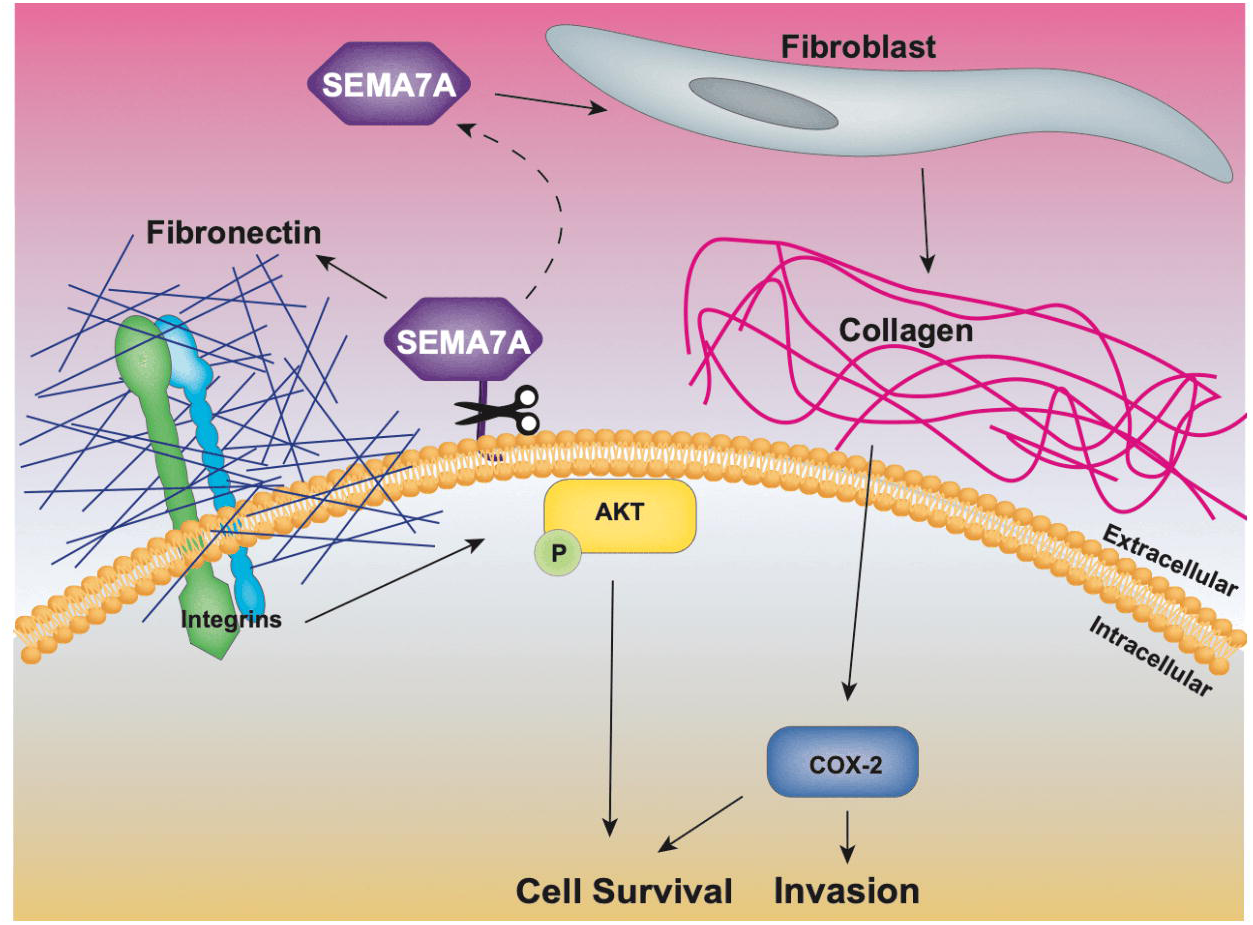
Model depicting SEMA7A mediated invasion and cell survival. Our current data suggests SEMA7A promotes fibroblast mediated collagen deposition, resulting in COX-2 expression and tumor cell invasion. Our results also suggest that SEMA7A can promote cell survival through SEMA7A mediated fibronectin expression.

SEMA7A was first identified on lymphocytes and originally designated CDw108 [33]. CDw108 was renamed SEMA7A due to its structural similarities with members of the Semaphorin family of proteins best known for their roles in neuronal guidance [34]. SEMA7A is the only semaphorin linked to the membrane via a GPI anchor; as such, it can be cleaved to result in shedding of SEMA7A into the extracellular environment, which was shown to promote β1-integrin dependent inflammation and fibrosis [35–41]. During fibrosis and in response to TGFβ, SEMA7A activates AKT signaling, which results in upregulation of collagen and FN in a β1-integrin dependent manner [42]. While our current studies do not explore a role for β1-integrin, we and others have shown that SEMA7A-β1-integrin binding promotes tumor growth, EMT, migration/invasion, metastasis, and neo-vasculogenesis [17, 19, 20]. Herein, we hypothesize that SEMA7A mediated tumor cell production of FN activates integrin mediated PI3K signaling leading to activation of AKT and cell survival [26, 43]. We also show that SEMA7A promotes mesenchymal phenotypes and tumor cell invasion, which could may be mediated by FN engagement of α_v_β1-integrin and downstream activation of Slug resulting in transcription of mesenchymal genes [44, 45]. In support of this hypothesis, SEMA7A is linked to EMT in a murine model of BC where TGFβ fails to induce EMT in the absence of SEMA7A [17]. Additional studies to determine how SEMA7A and FN contribute to EMT in our model are necessary to fully understand this mechanism in BC progression.

Since upregulation of EMT pathways such as Snail, Slug, and Zeb1 are associated with decreased sensitivity to chemotherapeutic drugs and targeted tyrosine kinase inhibitors (TKIs) [17, 46–55], it is also possible that SEMA7A may promote metastasis by conferring resistance to current therapies. In support of this, SEMA7A promotes resistance to EGFR targeted therapies in lung cancer [13]. Additional studies will explore the role of SEMA7A in resistance and/or susceptibility to current therapies in BC. Moreover, one major limitation of our current study is investigation of this mechanism only in ER-models, which is supported, in part, by our patient dataset studies showing that SEMA7A expression invokes a higher risk for metastasis in ER-breast cancers compared to all breast cancer. However, since ER+ breast cancers comprise the largest percentage of breast cancers in women, additional studies in ER+ breast cancers must be performed. Interestingly, estrogen and/or progesterone promote SEMA7A expression in the hypothalamus, suggesting that hormones may drive elevated levels of SEMA7A in BCs [56]. Furthermore, in hormone-receptor positive (HR+) BCs both AKT and FN promote tamoxifen resistance [57–64] and Semaphorin 4C, which is structurally similar to SEMA7A, drives hormonal-independence in HR+ BCs [65]. Thus, we are also exploring the relationship between SEMA7A expression and tumor progression in ER+ models [66].

We also demonstrate that SEMA7A is also involved in a non-cell-autonomous signaling axis via induction of fibroblast production of collagen, which then promotes COX-2 expression in the tumor cell [4]. Collagen has been characterized for multiple roles in promoting tumor progression as well as increased breast density, which can also increase risk for developing BC [8, 67, 68]. Additionally, a single pregnancy is sufficient to convert the high collagen content of the postpartum breast into pro-tumorigenic collagen [69] and activated fibroblasts, which deposit collagen, are present during postpartum involution [4, 8, 23]. Thus, our previous studies showing that SEMA7A is expressed in the postpartum mammary epithelium during involution SEMA7A, coupled with those presented herein, suggest that SEMA7A could also play a role in the increased risk for developing BC after pregnancy via increasing collagen deposition [1, 4, 31, 67, 68]. COX-2 is also well characterized for multiple roles in tumor initiation and progression [70, 71]. While our results suggest that SEMA7A stimulates collagen production by fibroblasts to result in tumor cell expression of COX-2 we still do not fully understand this mechanism. Additionally, our current studies have not addressed whether COX-2 activity is increased in tumor cells on collagen. Our data also suggest SEMA7A activation of AKT may play a role in promoting survival via upregulation of COX-2. Further studies are needed to understand the molecular mechanisms underlying SEMA7A and COX-2 signaling, regulation, and their roles in promoting tumor cell invasion and survival.

Currently there are no prevention strategies, targeted therapies, or specific treatment options for women diagnosed postpartum or for women with high tumor expression of SEMA7A. If our proposed SEMA7A mediated mechanism of progression is also dependent on COX-2 activity, as is suggested by our previous studies [4, 7], the addition of a COX-2 inhibitor to current treatment regimens could have efficacy in postpartum or SEMA7A+ BC patients. We also propose that SEMA7A expression could predict progression or itself be a potential therapeutic target for BC patients. If direct targeting of SEMA7A is not feasible, SEMA7A activates downstream targets such as FAK, Src, and ERK [16, 72], for which targeted therapies are available. Although these targeted inhibitors have been largely unsuccessful in BC, SEMA7A could serve as a predictive biomarker for patients who may benefit. Since metastases are the leading cause of BC related deaths and are largely untreatable, these novel anti-metastatic treatment strategies should be explored.

## MATERIALS AND METHODS

### Cell Culture

MCF10DCIS and MDA-MB-231 were cultured in 2D and 3D cultures as previously described [4, 18, 22]. MCF10DCIS cells and previously described shCOX-2 derivatives [73] were obtained from K. Polyak and A. Marusyk (Harvard University, Cambridge, MA). MDA-MB-231 cells were obtained from P. Schedin (Oregon Heath and Sciences University, Portland OR). HLF-1 cells were gifted from M. Fini (CU Anschutz Medical Campus, Denver, CO). Cells were validated by the DNA sequencing core at the CU Anschutz Medical Campus and identified to be a pure population of their respective cell lines. Cells were regularly tested for mycoplasma throughout studies. shRNA silencing was achieved using shRNA SEMA7A targeting plasmids (SABiosciences, Frederik, MD, and Functional Genomics Facility at CU Anschutz Medical Campus, Denver, CO) and confirmed via qPCR and Western blot analysis. Overexpression plasmid (SEMA7A-Fc) was a generous gift from R. Medzhitov (Yale University, New Haven, CT). Control plasmid (pcDNA3.1) was obtained from H. Ford (CU Anschutz Medical Campus, Denver, CO). All other overexpression plasmids (p304-V5-Blasticidin and V5-SEMA7A) were obtained from the Functional Genomics Core at the CU Anschutz Medical Campus and overexpression was confirmed via qPCR and Western blot analysis. Purified SEMA7A was isolated from MDA-MB-231 cells engineered to overexpress SEMA7A-Fc in collaboration with the Protein Purification/MoAB/Tissue culture core at the CU Anschutz Medical Campus. Cells were forced into suspension by coating plates with 12 mg/ml poly-HEMA (poly (2-hydroxyethyl methacrylate), Sigma, St. Louis, MO) prior to plating.

### Tissue microarray

Tissue microarrays containing normal and DCIS samples were prepared from biopsy tissue following placement in preservation media (LiforCell, Lifeblood Medical, Inc.) and storage at 4°C, as previously described [74].

### qPCR

RNA was isolated and qPCR were performed as previously described, with GAPDH and RPS18 as reference genes [18]. Primers for SEMA7A, COL1A1, COX-2 were obtained from Bio-Rad (Bio-Rad PrimePCR, Hercules, CA). Other primers were designed to be intron spanning with the following sequences: GAPDH (forward: CAAGAGCACAAGAGGAA GAGAG, reverse: CTACATGGCAACTGTGAGGAG) and RPS18 (forward: GCGAGTACTCAACACCAACA, reverse: GCTAGGACCTGGCTGTATTT).

### Immunoblot analysis

Western blots were performed as previously described [18]. Antibody information is provided in STable2.

### Animal model

The MCF10DCIS model was utilized as previously described [4, 18]. Briefly, 6-8-week-old female SCID Hairless Outbread mice from Charles River were bred and, after birth, pup numbers were normalized to 6-8 pups per dam. After 10-13 days of lactation, pups were removed to initiate involution (Day 0). Subsequently, injections of 250K MCF10DCIS controls and cells with shSEMA7A were initiated one day post-weaning (involution day 1) bilaterally into the #4 mammary fat pads. For SEMA7A overexpressing studies, 6-8-week-old Nude athymic (nulliparous) from Charles River were utilized because they are more cost-effective than SCID mice and breeding is not necessary for studies in nulliparous hosts. Tumors were measured twice weekly. For metastasis studies, nude mice were injected with 1x10^6^ cells into the tail vein, monitored for weight loss and sacrificed 3 weeks post-injection.

### Histologic analysis

Mammary glands with intact tumor were prepared for immunohistochemistry as previously described [4, 7]. Hematoxylin and eosin stained sections were examined by a board-certified anatomic pathologist (ACN) as scored as follows: 0-lesions which contained only well-devolved DCIS structures with clearly defined basement membranes and no evidence of microinvasion; 1-lesions that contained extensive DCIS with identifiable micro-invasive foci; 2-lesions that contained significant areas of sheet-like invasive tumor growth and mixed with areas of DCIS; 3-lesions that contained entirely invasive tumor with rare to absent DCIS remnants. The pathologist was blinded to study group by the randomization of animal numbers in each group.

### Immunohistochemistry and Immunofluorescence

For FN and COX-2 400X images were taken of intact tumor and quantitated using ImageJ software. For SEMA7A, cleaved caspase-3, and trichrome, stain quantification of total tumor area (necrotic and stromal areas removed) and percent positive stain or stain intensity was performed using ImageScope Aperio Analysis software (Leica, Buffalo Grove, IL). Areas for quantification were annotated using Aperio analysis tools and percent weak, medium, and strong determined using the color-deconvolution algorithm. For COX-2 and FN analysis, areas for quantification were isolated from the surrounding stroma and percent positive calculated as area with positive stain (positive pixels) using Image J and divided by total area (total pixels) and multiplied by 100.Immunofluorescent images were obtained using 400X magnification on OLYMPUS microscope. Antibody information is provided in STable 2.

### In vitro cell death assay

Cell death was analyzed using Caspase-Glo 3/7 Assay (Promega, Madison, WI), according to the manufacturer’s instructions.

### Analysis of publicly available datasets

Km plotter was queried for BC, and SEMA7A, PTGS2 (COX-2), and FN1 using the multigene classifier mean centered option for distant metastasis free survival (DMFS) [75]. ER-status was determined from ESR1 gene expression data. Ma breast cancer dataset was queried for SEMA7A, PTGS2 (COX-2), and FN1 using Gene expression-based Outcome for Breast Cancer Online (GOBO). TCGA was queried for co-expression of SEMA7A and COX-2 or FN1 using CBioPortal.

### Mass spectrometry analysis

Small tumor sections (∼1mm) were placed on gelatin sponges (Novartis Animal Heath, Greensboro, NC, USA) in serum-free media as previously described [76]. After 48 hours, tumor conditioned media was collected ∼ 30 µg of total protein digested utilizing the filter-aided sample digestion (FASP) protocol as previously described according to MAIPE standards [77]. Briefly, samples were reduced, alkylated, and enzymatically digested with trypsin. Resulting peptides were concentrated and de-salted by solid phase extraction utilizing in-house made stage tips made with Sytrene Divinyl Benzene disks (Empore™). Liquid chromatography tandem mass spectrometry (LC-MS/MS) was performed on a Thermo nanoEasy LC II coupled to a Q Exactive HF. MS acquisition parameters are detailed previously [78]. Raw files were searched with Proteome Discoverer 2.2 against the Mus Musculus, Homo Sapiens, and Bos Taurus uniprotKB database in Mascot. Precursor mass tolerance was set to +/-10 ppm and MS/MS fragment ion tolerance of +/-25 ppm. Trypsin specificity was selected allowing for 1 missed cleavage. Variable modifications include Met oxidation, proline hydroxylation, protein N-terminal acetylation, peptide N-terminal pyroglutamic acid formation, and a fixed modification of Cys carbamidomethylation.

Search results were visualized using Metaboanalyst v4.0 [79] and gene ontology mapping was done using PANTHER [80]. Data were prepared according to MIAPE standards and will be made available upon publication.

### Experimental replicates, sample size and statistical analyses

All in vitro studies were performed in biological triplicates. For animal studies, we chose the number of mice/group/time-point, based on power calculations form previous and pilot studies to achieve at least 80% power (β) with α=0.05. All animal studies were replicated twice with representative or pooled data shown. Unpaired and paired t-tests, ANOVA, and Kaplan Meier statistical analyses were performed in GraphPad Prism, assuming independent samples and normal distributions. Analyses for Figure 2 were done using one-tailed t-tests, as our results from Figure 1 would predict significant differences between groups. All other analyses were done using two-tailed tests. Only p-values of less than 0.05 were considered significant. Aside from unexpended death or cell contamination, no samples or animals were excluded from our analysis. For IHC analysis of tumors, outliers were removed if they were significant by the ROUT (Q=1%) test.

### Study Approval

Prior to resection, patients provided written informed consent under an IRB-approved protocol according to the guidelines of their respective institutions and conducted in compliance with HIPPA regulations. All animal studies were approved by the IACUC of the CU Anschutz Medical Campus, protocol number B106017(06)1E.

## Author Contributions

SET and TRL conceived and designed the study. SET performed all in vitro and in vivo studies. VFB and FB were responsible for regulatory oversight of human tissue acquisition and providing cases for IHC analysis. RCH and KH were responsible for all mass spectrometry experiments and associated data analysis. ACN was responsible for analyzing and scoring all tumor for invasion. SET and TRL were responsible for hypothesis development, conceptual design, data analysis and data interpretation. SET and TRL wrote the manuscript with all authors providing critical evaluation.

## Acknowledgements

MCF10DCIS cells were obtained from K. Polyak and A. Marusyk (Harvard University, Cambridge, MA). MDA-MB-231 cells were obtained from P. Schedin (Oregon Heath and Sciences University, Portland OR). HLF-1 cells were gifted from M. Fini (CU Anschutz Medical Campus, Denver, CO). H. Ford for the pcDNA3.1 vector (CU Anschutz Medical Campus, Denver, CO).; and R. Medzhitov (Yale University, New Haven, CT) for the SEMA7A-Fc overexpression vector. We also acknowledge V. Wessells, A. Elder, L. Crump, T. Wallace, C. Young, A. Stoller, M. Kobritz, and C. Hoang for technical support and advice. This work was supported by the American Cancer Society (RSG 16-171-01-CSM), and NIH/NCI (R01CA211696-01A1) to T. Lyons; NIH/CCTSI/CTSA (TL1 TR001081) to S. Tarullo.

**Supplementary Information** is available on *Oncogene’s* website.

