## Supplemental Data for "Postpartum breast cancer progression is driven by semaphorin 7a mediated invasion and survival"

### SUPPLEMENTAL FIGURES

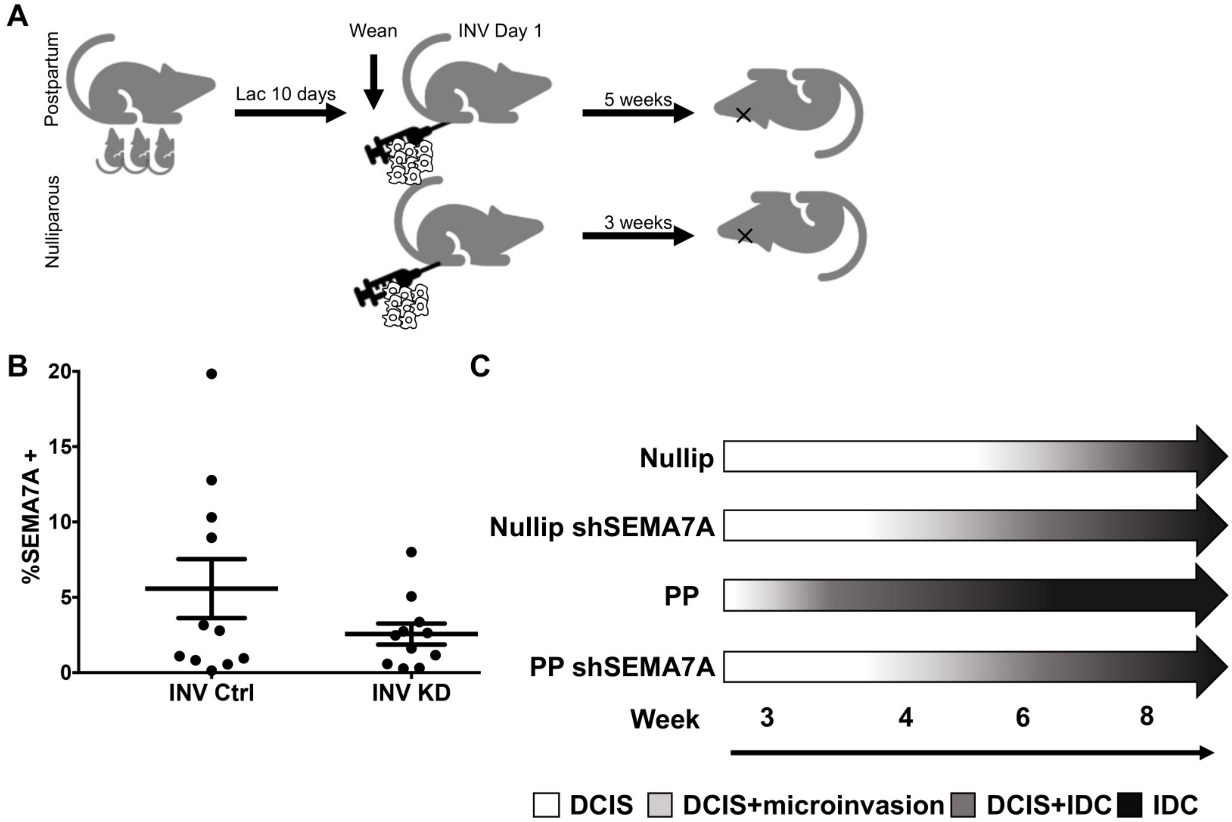


**SFigure 1. A.** Schematic of *in vivo* model. **B.** Quantification of SEMA7A stain in control (INV Ctrl) and SEMA7A KD (INV KD) MCF10DCIS tumors from postpartum hosts. **C.** Diagram summarizing effects of SEMA7A knockdown and parity status on MCF10DCIS xenograft tumor progression.

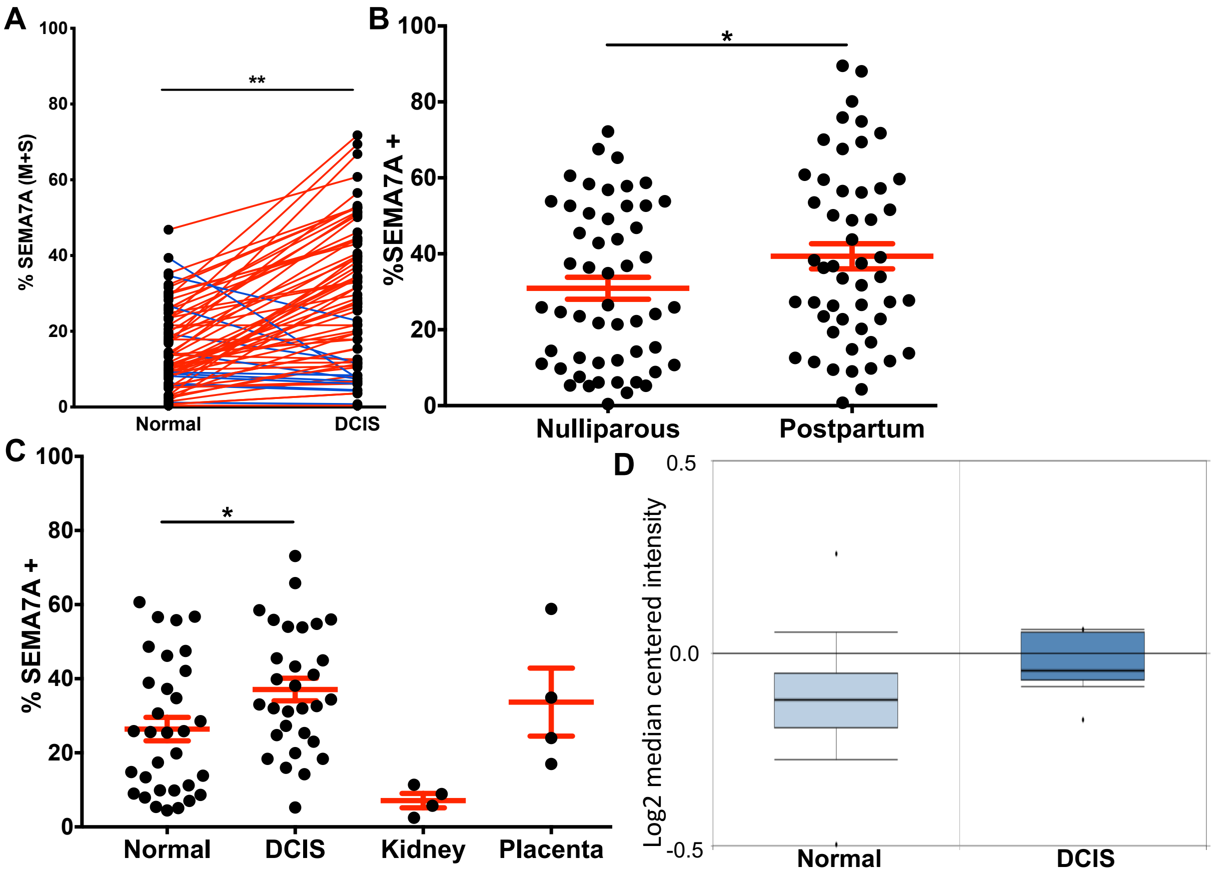


**SFigure 2. A.** Quantification of SEMA7A in normal or DCIS tissue from YWBC cohort by patient, paired t-test. **B.** Quantification of SEMA7A in DCIS tissue from nulliparous or postpartum patients. **C.** Quantification of SEMA7A stain from tissue microarray in normal human breast tissue, all breast cancer patients with DCIS, kidney (negative control), or placenta (positive control). **D.** Oncomine analysis for gene expression of SEMA7A in normal versus DCIS in the METABRIC dataset (p=0.002). (*p<0.05, ****p<0.001).

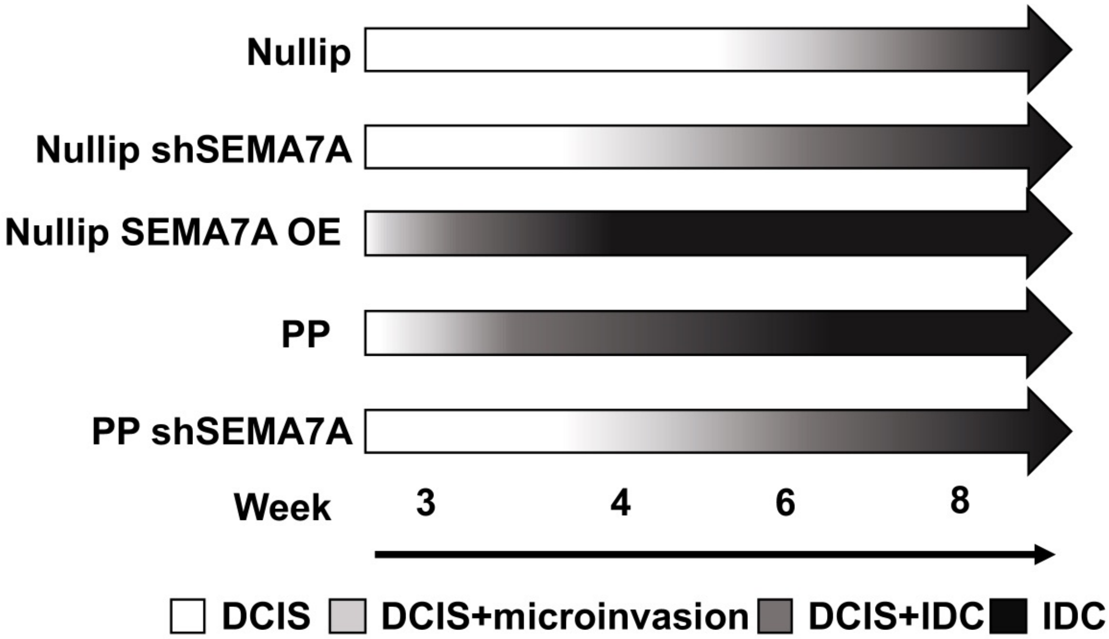


**SFigure 3.** Diagram summarizing effects of SEMA7A expression and parity status on MCF10DCIS xenograft tumor progression.

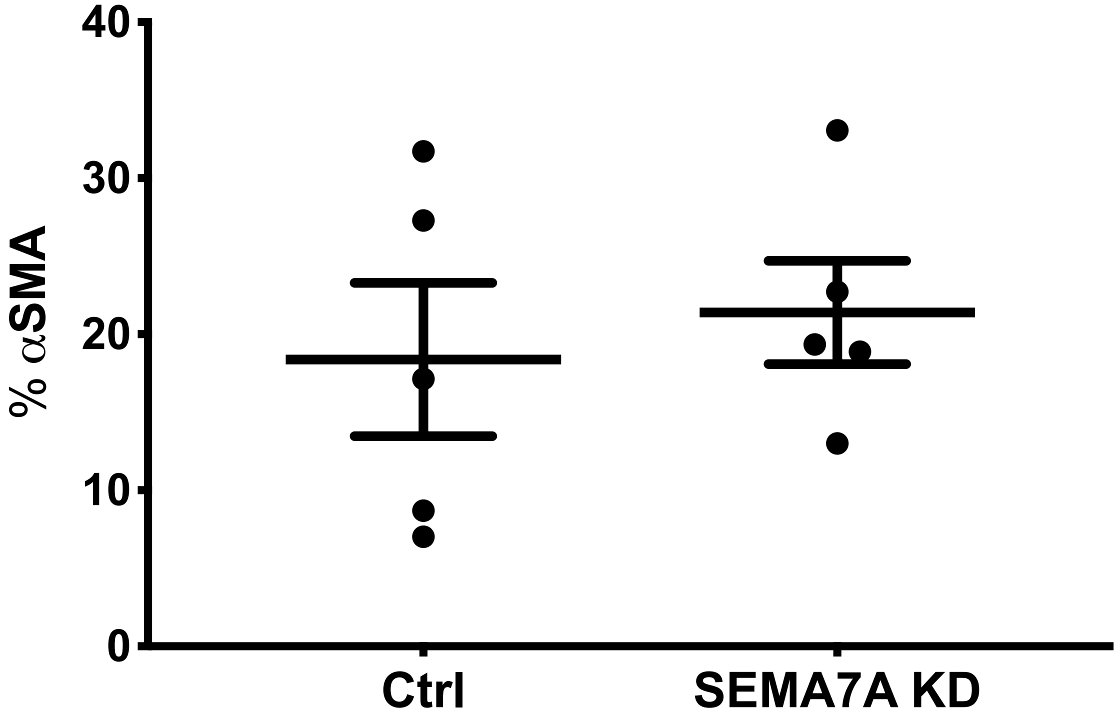


**SFigure 4.** Quantification of α-smooth muscle actin (αSMA) of Ctrl and SEMA7A KD tumors from postpartum hosts.

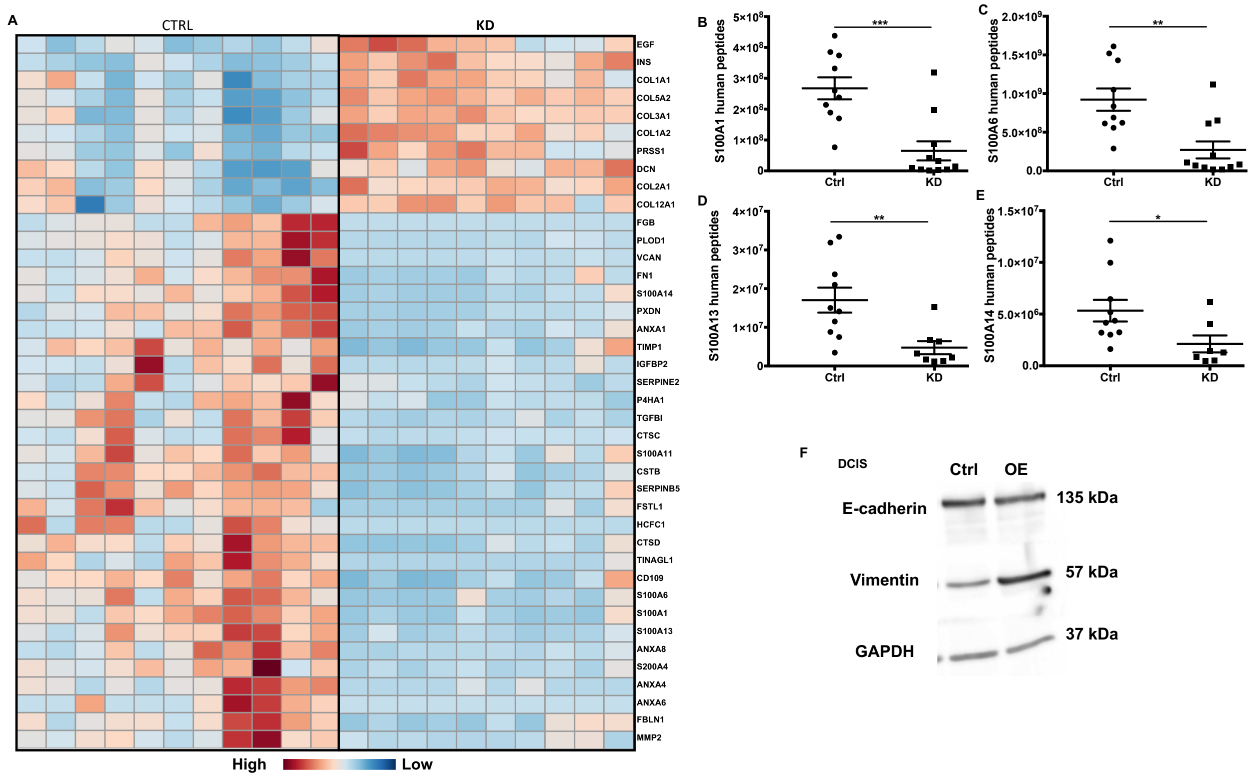


**SFigure 5. A.** Clustering analysis of human extracellular matrix proteins following global mass spec analysis. **B-E**. Additional decreases in mesenchymal or mesenchymal-associated proteins observed by proteomics in SEMA7A KD tumors *ex vivo*.

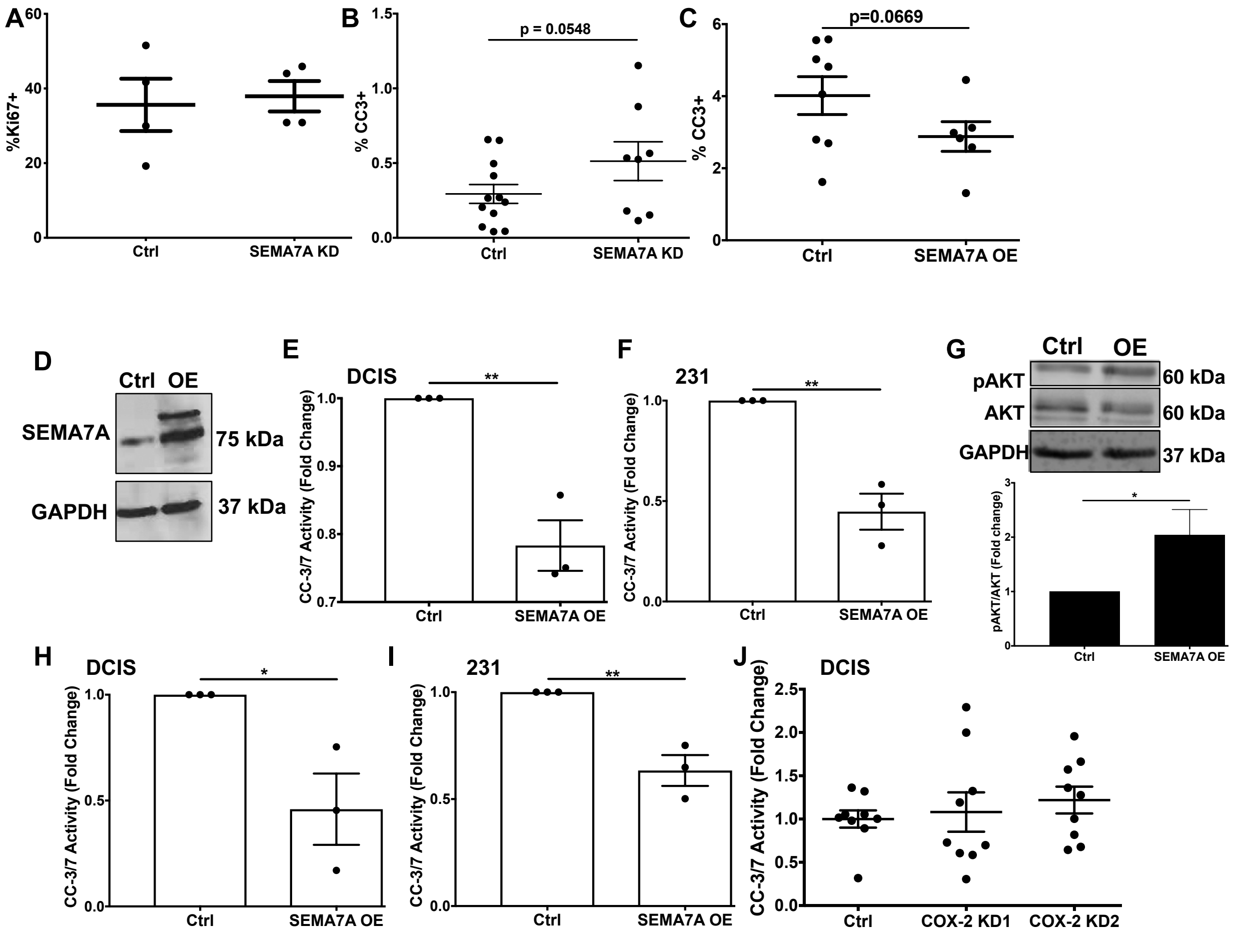


**SFigure 6. A**. Quantification of Ki67 of Ctrl and SEMA7A KD tumors from postpartum hosts. **B&C**. Quantification of cleaved caspase-3 (CC3) of Ctrl and SEMA7A KD from postpartum hosts or control (NUL Ctrl) or SEMA7A overexpressing (NUL SEMA7A OE) tumors from nulliparous hosts. **D.** Representative immunoblot for SEMA7A in Ctrl or SEMA7A OE MDA-MB-231 cells. **E&F.** Fold change of cleaved caspase 3/7 activity in control (Ctrl) and SEMA7A overexpression (SEMA7A OE) cells in MCF10DCIS (**E**) or MDA-MB-231 (**F**) cell lines. **G.** Representative immunoblot for phospho-AKT (S473), total AKT, or GAPDH in Ctrl and SEMA7A OE, quantified below. **H&I.** Fold change of cleaved caspase 3/7 activity in control (Ctrl) and SEMA7A overexpression (SEMA7A OE) cells in MCF10DCIS (**H**) or MDA-MB-231 (**I**) cell lines in forced suspension. **J.** Fold change of cleaved caspase 3/7 activity in Ctrl or shCOX-2 (KD1/KD2) cell lines plated on tissue culture plastic.

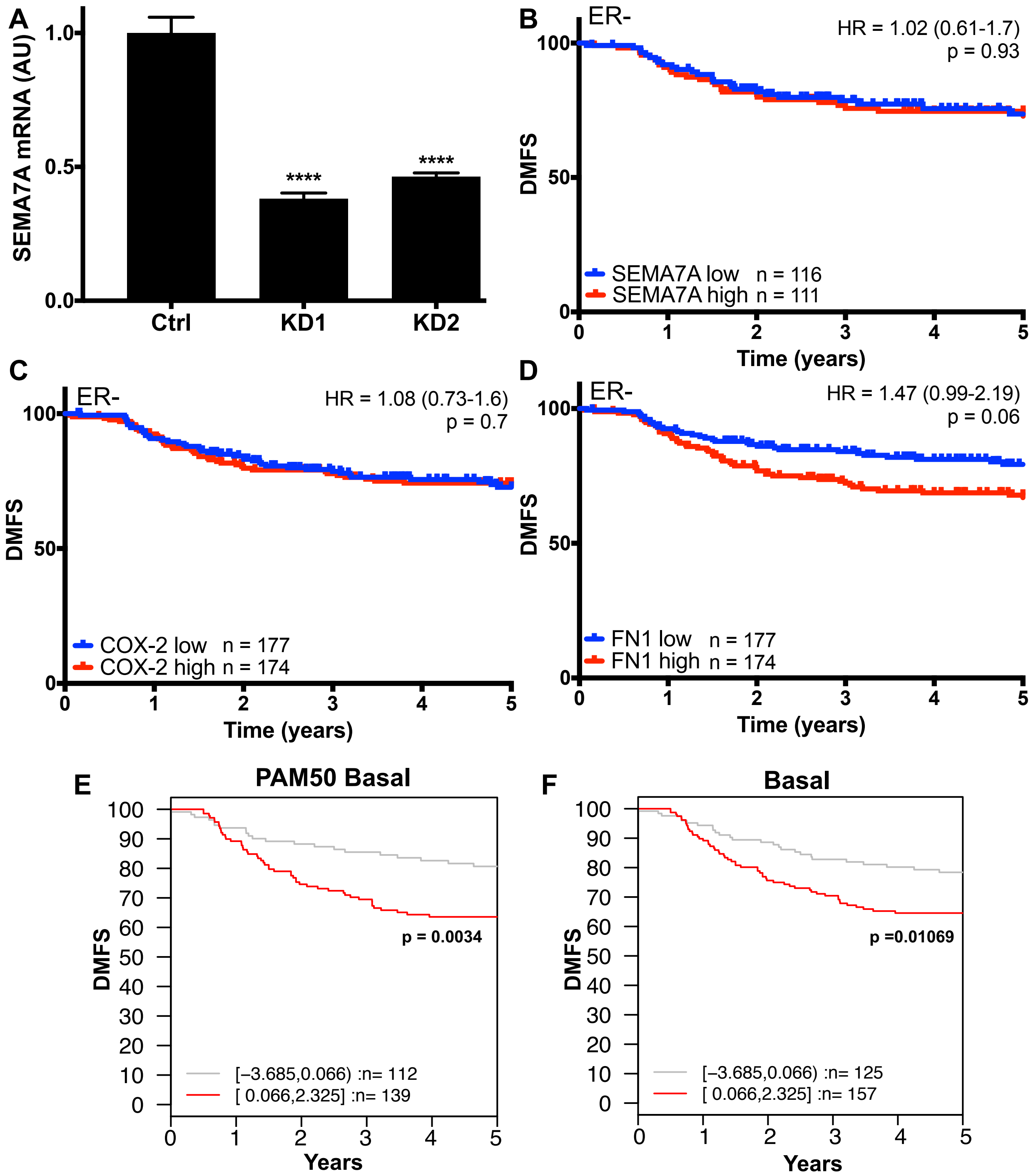


**SFigure 7. A**. Representative qPCR for SEMA7A normalized to GAPDH and RPS18 in MDA-MB-231 cells used for tail vein injections. **B.** Kaplan-Meyer analysis of estrogen receptor negative (ER-) breast cancer patient samples using Km plotter for SEMA7A for distant metastasis free survival (DMFS) in. **C&D.** Kaplan-Meyer analysis of ER- breast cancer patient samples using Km plotter for (**C**) COX-2 and (**D**) FN for DMFS. **E&F.** Kaplan-Meyer analysis Ma Breast Cancer dataset for DMFS in PAM50 Basal (**E)**, and basal (**F**) breast cancers using GOBO. (****p<0.001).

**STable 1.** Table summarizing expression of SEMA7A, COX-2, and fibronectin (FN) in normal versus cancer patient samples, data obtained from Km plotter.

| mRNA | Tissue | Median | P value | Sample n |
|---|---|---|---|---|
| SEMA7A | Normal | 97 | NA | 76 |
| | Cancer | 112 | 0.00697 | 6547 |
| COX-2 | Normal | 309 | NA | 76 |
| | Cancer | 71 | 3.87e-15 | 6547 |
| FN1 | Normal | 4193 | NA | 76 |
| | Cancer | 17920 | 1.28e-28 | 6547 |

**STable 2.** Antibodies used for immunoblot, immunohistochemistry, and immunofluorescence.

| Antibodies | Clone | Manufacturer | Dilution | Antigen Retrieval |
|---|---|---|---|---|
| Semaphorin 7A | C6-Human tissues and WB D4-xenograft | Santa Cruz Biotechnologies | 1:500 (IHC) 1:250 (WB) | TRS (pH 6.0) |
| Cleaved Caspase-3 | Asp175 | Cell Signaling Technologies | 1:100 | EDTA (pH 9.0) |
| Ki67 | SP6 | Thermofischer Scientific | 1:400 | TRS (pH 6.0) |
| COX-2 | N/A | Cayman Chemical | 1:250 | TRS (pH 6.0) |
| $\alpha$SMA | N/A | Abcam | 1:200 | TRS (pH 6.0) |
| Fibronectin | N/A | BD Biosciences | 1:2000 | EDTA (pH 9.0) |
| F-actin Alexa Fluor 568 phalloidin | N/A | Invitrogen | 1:50 | N/A |
| pAKT (S473) | D9E | Cell Signaling Technologies | 1:2000 | N/A |
| AKT | 40D4 | Cell Signaling Technologies | 1:2000 | N/A |
| Goat anti-mouse secondary | N/A | Abcam | 1:5000 | N/A |
| Donkey anti-mouse | N/A | LiCOR | 1:15,000 | N/A |
| Donkey anti-rabbit | N/A | LiCOR | 1:20,000 | N/A |
